# Insights into the molecular basis of wooden breast based on comparative analysis of fast- and slow-growth broilers

**DOI:** 10.1101/356683

**Authors:** Shawna Marie Hubert, Travis J. Williams, Giridhar Athrey

## Abstract

Wooden breast has emerged as an important condition in the poultry industry and affects the breast muscle of commercial meat-type chickens (broilers). Thus far, the condition has been classified as a myopathy, confirmed to not have an infectious origin, with molecular data showing the muscle to be under oxidative stress. The objective in this study was to query the functional origins of wooden breast and reveal its molecular similarities to known conditions based on pathway analyses. To carry out an in-depth comparative analysis, we generated RNAseq data from wooden breast affected birds and incorporated breast specific transcriptomic data from previously published studies. The comparative datasets were constructed from a range of commercial fast-growth and slow-growth varieties. Analysis of high-impact variants identified from transcriptome data provided a list of genes important in cell signaling, cell proliferation, cytoskeletal development, and calcium metabolism as being affected by nucleotide changes. Overlaying the lists of significantly differentially expressed genes with the list of high-impact variants produced a list of twenty genes that suggest an association of mechanistic and functional causes for woody breast; supporting a polygenic basis for this condition. Our study demonstrates that wooden breast shows an age-dependent gene expression pattern, with pathway analysis showing enrichment of glycolysis, cell differentiation, tumor suppression and inhibition, further indicating a complex condition with few similarities to myopathies. In summary, our results indicate the existence of a mechanistic, heritable basis for wooden breast, the drivers of which deserve more in-depth investigation. Additionally, they also suggest wooden breast to be a more complex condition than previously reported, potentially involving other organ systems.

## Introduction

The domestic chicken (*Gallus gallus domesticus*) is a major agricultural species and is arguably the most popular source of animal protein around the world. In the United States, chicken breast is the most consumed meat – per capita consumption surpassed 41kgs in 2015 (source: US Poultry) – and the broiler industry has a substantial economic footprint ($30 billion/year). Consumption of poultry has increased in step with human population growth as well as changes in consumption habits [1]. While demand continues to grow, production is under enormous stress due to a variety of disorders (ascites, fatty liver disease) and meat quality issues such as green muscle disease and wooden breast [2–6]. Of these, wooden breast (WB) is the most recent problem that is negatively impacting breast meat quality. WB is a muscle condition categorized as a breast myopathy, causing increased hardness of tissue and reducing meat quality. The frequency of WB has risen steadily over the last five years, being reported globally with reduced consumer acceptability [7] linked to economic losses [8, 9]. WB has become prominent within the last decade and has been reported to affect over 50% of commercial flocks [10, 11], but accurate estimates of global incidence are not known.

Pectoral myopathies are not new in broiler poultry species, and broiler chicken particularly has a well-documented history of dystrophies and myopathies, including pectoral myopathies induced by physical or nutritional stress [12–14]. For example, Siller *et al.* [15] reported deep pectoral myopathy in both turkeys and broiler chicken induced by exercise.

However, WB is different from previously studied pectoral myopathies in some meaningful ways; the hallmarks of WB appear to be moderate to severe degenerative necrosis, with varying degrees of interstitial fibrosis. While some of these features are found in other myopathies, the co-occurrence of localized pectoral myopathy with fibrosis and striations has not been observed previously in broilers. WB is often observed in conjunction with white striping (WS), which is characterized by white striations that run parallel to the muscle fibers in the breast [11]. These white striations can resemble marbling and are associated with increased fat content [16]. In the past, some pectoral myopathies have been assigned etiologies ranging from nutritional deficiency (e.g., selenium) to hypoxia or ionophore toxicity [14, 17]. However, these etiologies are not supported in WB. Despite intensive studies of WB, including histopathological analyses, serological studies, dietary interventions, gene expression, and metabolomics studies [10, 18–22], the causative factors of WB remain unknown.

Both WB and WS demonstrate varying degrees of severity and have been identified in multiple commercial varieties. Recent studies have described WB as a polyphasic myodegeneration [11], presenting lymphocytic phlebitis [11, 20]. The only common explanatory factor appears to be the **growth rate of commercial broilers**, with the most severe cases found in the heaviest male birds [19, 23]. Kuttappan *et al.* [3] reported that the rapid growth rate and high-energy diets both increase the incidence of WS. A similar trend is observed with WB. Since 2000, average broiler weights have increased by 3 kg (6.5 lbs), representing a 55% increase (Source: U.S. Poultry). Due to the high value of breast meat in proportion to the total carcass, increasing incidence and severity of WB translates into more significant economic losses [24].

While selective breeding for performance traits (e.g. growth rate, feed efficiency) and advances in nutrition, are primarily responsible for growth rate improvements in broilers, it is not known whether WB, which is associated with growth rate, has a genetic basis. Recent studies have used gene expression (RNAseq) and metabolomics analyses to characterize WB [9, 10, 18], but these investigations have not been informative about the underlying cause(s) of WB. Whereas a previous report suggested low heritability for WB [24], a recent report by Pampouille *et al.* [25] describes the identification of quantitative traits loci (QTL) for WS in high-yield broilers, and further concludes that WS is a polygenic condition, supporting the hypothesis for a strong genetic basis for WB and WS.

In this study, the objective was to utilize comparative analyses to illuminate the basis of WB and to characterize its similarity to other known conditions. This study also evaluated the evidence for the supposition that WB is a muscle myopathy. We addressed this objective by comparative transcriptomic analyses of WB samples against various genotypes/phenotypes, followed by pathway analyses and tests for enrichment of canonical pathways. This study did not specifically focus on molecular features of WB/WS co-occurrence, and hence we do not draw inferences regarding WS. Altogether, our study indicates that a) WB is an age-dependent disorder driven by transcriptional dysregulation in fast-growth broilers, and b) that WB molecular profiles suggest a complex syndrome potentially involving multiple organ systems. These results suggest a genetic basis to WB and emphasize the importance of more in-depth studies of the mechanistic basis of WB. These findings also suggest that WB is a condition with potential consequences for whole organism health.

## Materials and Methods

### Study design and source of data

In this study, transcriptome data was generated from birds exhibiting WB, which was then analyzed comparatively with data generated for previous broiler gene expression studies and published in the peer-reviewed literature. The details of these samples, with links to original studies, are provided in Table 1. It is important to note that these studies focused on breast tissue-specific gene expression and to our knowledge, WB was not the explicitly stated subject of investigation. However, a subset of these studies used modern commercial broilers. Therefore we cannot be sure that they were unaffected by WB; however based on the high frequency of WB in commercial broiler flocks [21] it is likely that these samples include WB affected individuals.

**Table 1:** Summary of data used in the comparative analysis, including information about the chicken breed, tissue type, the age of birds at sampling, SRA accession info, and authors of the original study.

| SRA<br>accessio<br>n ID | Chicken<br>Breed/variety | Breed Type | Tissue<br>Type | Age of bird<br>at sampling | Authors |
| --- | --- | --- | --- | --- | --- |
| PRJNA3<br>39392 | Arbor Acres | Commercial Broiler | Pectorali<br>s major | 42-days | Yi et al.,<br>2016 [81] |
| PRJNA3<br>42997 | WC and WRR | Heritage Broilers | Pectorali<br>s major | 120 and 180<br>days | Qui et al.,<br>2017 [82] |
| PRJNA2<br>94010 | Recessive White<br>Rock (WRR) &<br>Xinhua | Heritage &<br>Indigenous | Pectorali<br>s major | 42-days | Chen et al.<br>[83] |
| PRJNA2<br>73416 | Ross 708 & Illinois<br>Chickens | Commercial Broiler<br>& Legacy Breed | Pectorali<br>s major | 6 and 21-days | Davis et<br>al., 2015<br>[84] |
| PRJNA2<br>66323 | WRR & XH | Heritage &<br>Indigenous | Pectorali<br>s major | 42-days | Ouyang et<br>al., 2015<br>[85] |

WRR: White Recessive Rock, WC: Wenchang Chicken, XH: Xinghua Chicken

To determine the common genetic basis for WB as a distinct signal, aside from the breed specific growth and molecular profiles, it is necessary to compare expression profiles across ages and genetic strains. WB has been reported in fast-growth broilers as early as two weeks of age, but with the most dramatic changes in severity occurring in the final three weeks before slaughter [16, 22]. Therefore, we compared gene expression of pectoralis major muscle tissue from fast- and slow-growth broilers. While WB has been reported from all major commercial broiler strains, WB has not been reported in slow-growth and heritage broilers to date.

Secondly, as WB severity has been reported to increase with age and weight, we compared expression profiles among pectoralis muscle tissues from different age categories of both fast- and slow growth. To answer these questions, we used a combination of data generated in-house (reported above) in addition to reusing publicly available sequence data (NCBI Short Read Archive) that matched our analysis criteria. In total, nine publicly available datasets from six previously published studies were included in these comparisons (Table 1). In each instance, we selected studies that generated RNAseq data from the breast tissue, were sequenced on the Illumina platform, and were not from a pathogen challenge experiment. Three datasets were from an environmental ammonia challenge study, using the 42-day old broilers of the Arbor Acres strain (Aviagen, sample prefix ARAC). While these treatments influence their gene expression profiles, our analyses showed that these three sample groups cluster together with other 42-day old fast-growth broilers. The inclusion of data from this experiment did not change the hierarchical clustering and separation of sample transcriptome profiles by performance and age profile; hence we retained all three datasets in further analyses.

### Sample Collection and RNA Extraction

Live animal studies and euthanasia procedures performed in-house were approved by the Texas A&M University’s Animal Care and Use Committee (assurance number 2016-0065). Breast tissue samples were collected from eight, 42-day old chickens of a high-yielding commercial broiler strain. The birds in the study were from an all-male flock and raised on a three-phase industry standard diet. Two birds were randomly sampled from each of four replicate pens, containing 40 birds each. The breast muscle was palpated to examine birds for hardness, before euthanasia. This approach has been used as a diagnostic method in several recently published studies [10, 26, 27]. Birds were then euthanized by cervical dislocation and dissected for the collection of tissues for genetic analysis. The pectoralis major and pectoralis minor muscles were then examined for gross lesions and hardness of the muscle. Individual samples were classified as either WB+ or WB- based on the observed hardness of breast tissue and the absence of other visible abnormalities. While histological analyses have also been used for WB classification, they are perhaps more applicable for resolution of severity, rather than a diagnostic for presence-absence of WB. Furthermore, histological classification of breast tissue as ‘normal’ has not been found to be diagnostic of WB at the molecular level [11, 28]. While it has been noted that WB and WS co-occur frequently, this study was focused on WB, and therefore did not specifically classify tissue for WS presence or severity. Of the eight individual birds sampled this way, six birds were classified as severe (WB+) based on palpation and gross lesions, whereas two other samples were less severe cases (WB-). Owing to the weak correlation between physical/histological and molecular markers of WB, all birds sampled in this study were classified into the WB group. Moreover, as the goal was to compare expression patterns across genotypic backgrounds, this grouping allowed better resolution through increased biological replication of the WB group.

Tissue of size approximately 1cm^3^ was excised from the distal portion of the pectoralis major with a scalpel and immediately stored in RNAlater (Ambion Inc). After 24 hours of incubation at 4°C, the excess RNAlater was removed, and samples were stored at −80°C until further processing. Total RNA was extracted from about 30mg of the tissue using the RNEasy Mini Kit (Qiagen Inc). Samples were checked for RNA quality and concentration on a NanoDrop ND-1000 Spectrophotometer (Thermo Fisher Scientific).

### RNA Sequencing and Transcriptome Analysis

Total RNA isolates were submitted for library preparation and RNA-sequencing at the AgriLife Genomics and Bioinformatics Center (Texas A&M University). Sample libraries were prepared by performing DNAse digest, followed by poly-A selection for mRNA molecules. Individual mRNA isolates were then pooled into three sample libraries – namely one library comprising two WB- samples, and two libraries each comprising three WB+ samples. These three sets of pooled samples were used for strand-specific library preparation, and the libraries were sequenced with 125bp single end sequencing.

The 11 datasets, including the in-house generated and downloaded datasets, were then processed identically. In brief, the raw RNAseq data was filtered for adapter contamination and quality trimmed using the program Trimmomatic [29]. Reads with average quality scores less than Q30 and shorter than 20bp in length were discarded. High-quality reads were mapped to the *Gallus gallus* genome (Version 4.8, Ensembl Release 85, July 2016) using the short-read de-novo splice mapper STAR [30, 31], followed by counting of transcripts mapped to the ‘exon’ features using the tool HTSeq [32]. The counts data for each sample were then compared for statistical significance using the EdgeR package on the R statistical platform [33]. First, low expressed genes across all libraries (CPM <2) were filtered out. Next, normalization factors were calculated for differences in library sizes, followed by estimation of common and then tagwise dispersion (GLM). We used the package COMBAT to check and correct for batch effects [34, 35]. The quasi-likelihood based ‘glmQLFTest’ function was used to perform tests for significance between expression values among treatments. The QLF approach is known to provide greater protection against Type I error and can handle unbalanced designs better than exact tests. A total of 10 pairwise contrasts were performed between WB data (generated in-house) and downloaded broiler transcriptome data. Following the analyses in EdgeR, topTag tables were used in interpretation and pathway analyses. For each of the ten differential expression results, pathway analyses were performed using the Ingenuity Pathway Analysis platform (Qiagen Inc.). Only genes significant at FDR <0.05, and with Log2FoldChange smaller than −0.5 or greater than 0.5 were included in pathway analyses. Finally, the results from pathway core analyses (10 datasets) were all included in a ‘Comparison Analyses’ on the IPA platform, to characterize similarity of expression, shared canonical pathways, upstream regulators, and diseases.

### Differential expression between Fast-Growth versus Slow-Growth broilers

A secondary differential gene expression analysis was performed by grouping all the modern commercial broilers as the Fast-Growth Commercial Broiler (FGCB), and by grouping the Weichang (WC) and White Recessive Rock (WRR) samples as the Slow-Growth Heritage Broiler (SGHB). The Illinois strain and the hybrid WRR-XH crosses were left out of this comparison as they are neither a heritage breed nor a commercial variety. Differential expression analysis and pathway analysis was performed in the same way as described above.

### Variant analysis with RNAseq data

A total of 54 sequence libraries (.fastq), including the eight generated for this study, were used to generate variant calls and to identify shared and unique SNP variants among the breeds included. The Genome Analysis Toolkit [36] best practices pipeline for variant calling from RNAseq data was used to generate a set of high-quality variants for each sample using hard filtering. Briefly, STAR aligned reads (same used for differential expression analysis) were first processed to add read information and to mark duplicates using the tool Picard [37]. Binary alignment files were then fed into HaplotypeCaller, followed by the selection of SNP variants, and variant filtration. SNPs occurring in clusters within 35bp were filtered out, as were variant calls with Qual By Depth (QD) score <5, and Fisher strand bias > 35. The resultant set of high-quality variants obtained this way were passed into the variant effect prediction software SnpEff [38]. Variants annotated as having a ‘High’ impact modifier by SnpEff were used for comparison among samples. Due to the variability in sequencing library size and depth of coverage, differences in the number of variants were expected. Therefore, to account for potential bias arising from differences in sequencing depth, high-impact variants were compared only between the FGCB group, and the SGHB group. The combined high-impact variant list was generated by pooling all variants by ENSEMBL gene ID and removing duplicates.

## Results

### Growth rate and age explain global gene expression patterns

Across the ten datasets, a total of 12,202 genes were expressed above threshold (CPM=>2) and were included in both the differential expression and pathway analyses. An ordination analysis using Non-metric Multidimensional Scaling (NMDS, Figure 1) showed that fast-growth breeds (ARAC, ROSS and WR-XH Cross) overlapped each other, with younger (6 and 21-days old) fast-growth broilers being less proximate to WB samples, compared to 42-day old broilers (ARAC), indicating a clear age based expression similarity. The Illinois and Ross breeds (6 and 21-days old) both clustered by age and also by breed, showing clear age-based segregation from 42-day old commercial broilers. The slow-growth breeds (WC and WRR, 120 days and older) formed clusters distinct from the fast-growth broilers. Furthermore, all WB samples formed a tight cluster, validating the observation that birds without obviously visible WB symptoms are, nonetheless, not different at the molecular level. Therefore, birds from the same genetic background may not be suitable as a negative control (Figure 2).

**Figure 1:**
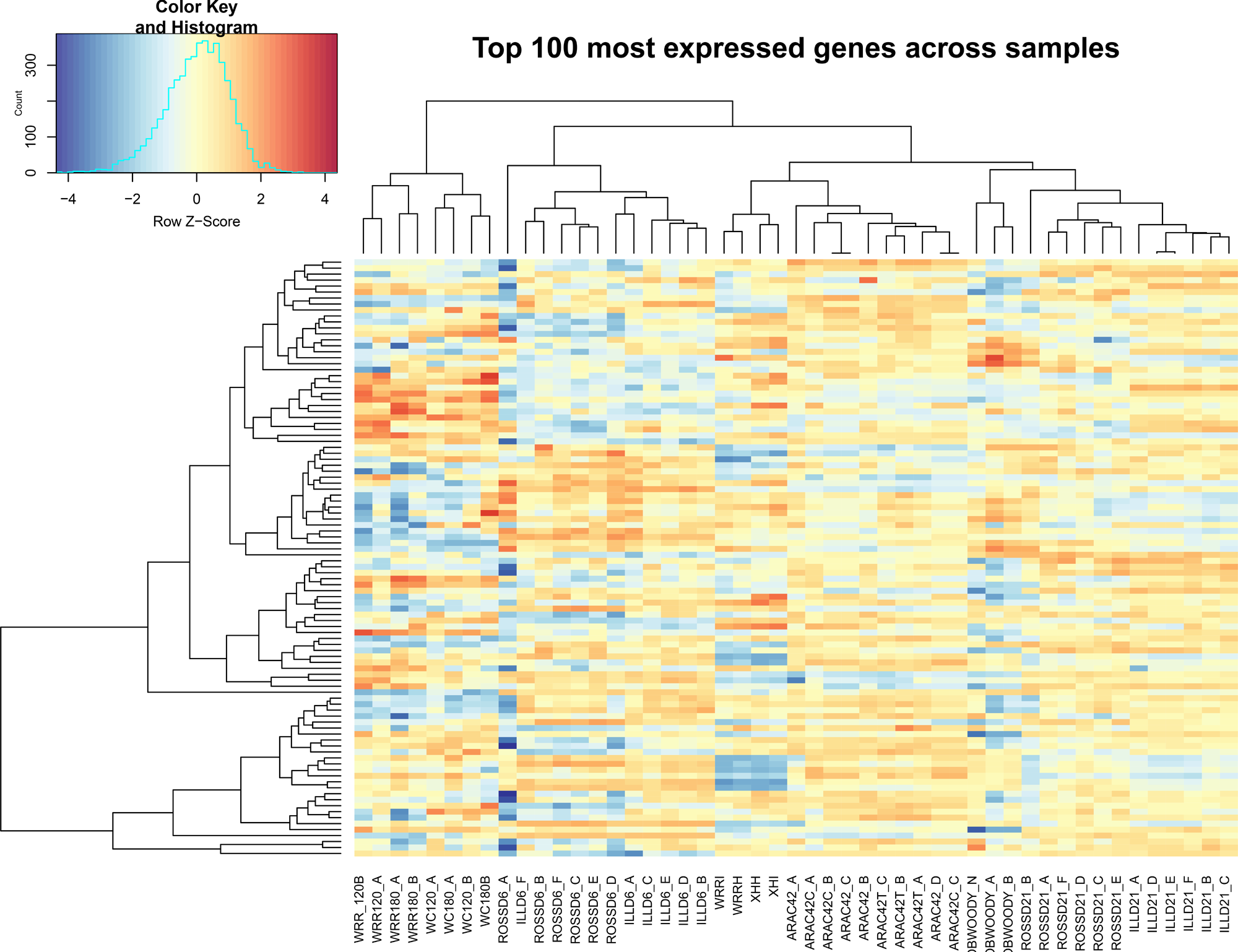
Non-Metric Multidimensional Scaling plot of the RNAseq datasets in the multi-sample comparison study. The samples from each age and growth-rate group cluster clearly within groups. Commercial fast-growth broilers also appear more proximate to each other and are arranged from top to bottom in order of increasing age, while youngest and slowest growth breeds are furthest away from older, fast-growth breeds.

**Figure 2:**
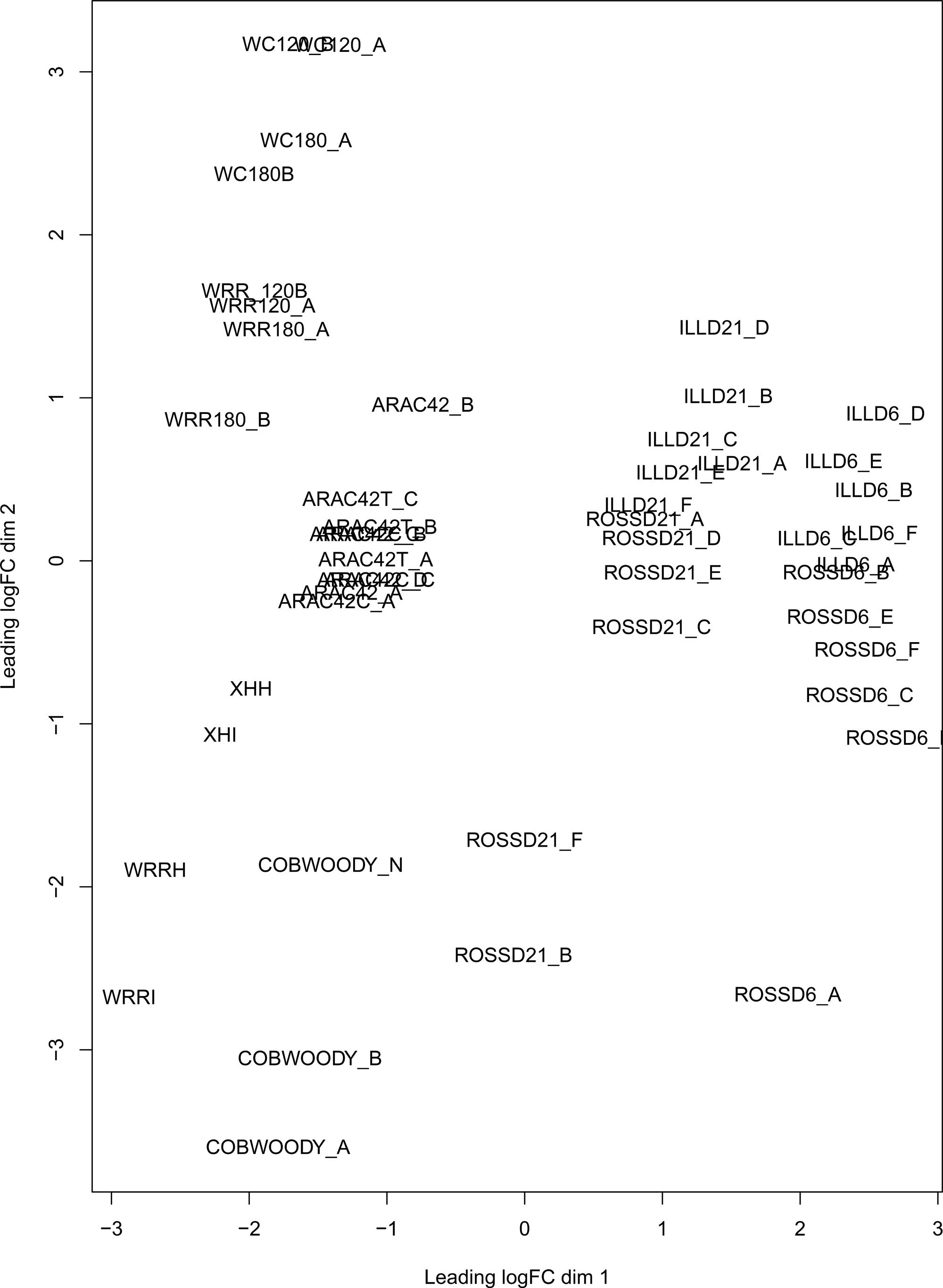
Hierarchical clustering heatmap of the top 100 most expressed genes, based on logCPM values, across samples. Clustering shows that woody breast samples fall among Ross 21-day old and ARAC 42 old birds, with slow-growth heritage birds (WC and WRR) forming a distinct cluster.

### Comparison of pairwise differential expression analyses

Results from EdgeR for each pairwise comparison against WB samples are summarized in Table 2, and mean-average plots for each comparison is shown in Figure 3. The complete list of differentially expressed genes with P-values for all pairwise comparisons is available in supplementary materials. Overall, the WC120+ and WRR120+ slow-growth varieties were most different from WB, based on the total number of differentially expressed genes (~2200). This number was approximately twice as high as any of the other comparisons. After the WC and WRR breeds, the Illinois 21D was most different (1702 genes differentially expressed), and Ross 21D being most similar (693 genes differentially expressed), with other comparisons falling in between the extremes. The three different datasets of 42-day ARAC (3 treatments in the original study), were very similar to each other in their differences to WB (total of 1243, 1167, and 1330 DEG respectively). The top canonical pathways identified were also highly similar, with T-cell receptor signaling in all three comparisons and IL-8 signaling in comparison to both A and C groups of ARAC. Canonical pathways explained by these DE genes showed that IL-8 signaling and T-cell receptor signaling were recurrent terms, but all three comparisons shared TGFB1 and TNF as the upstream regulators.

**Figure 3:**
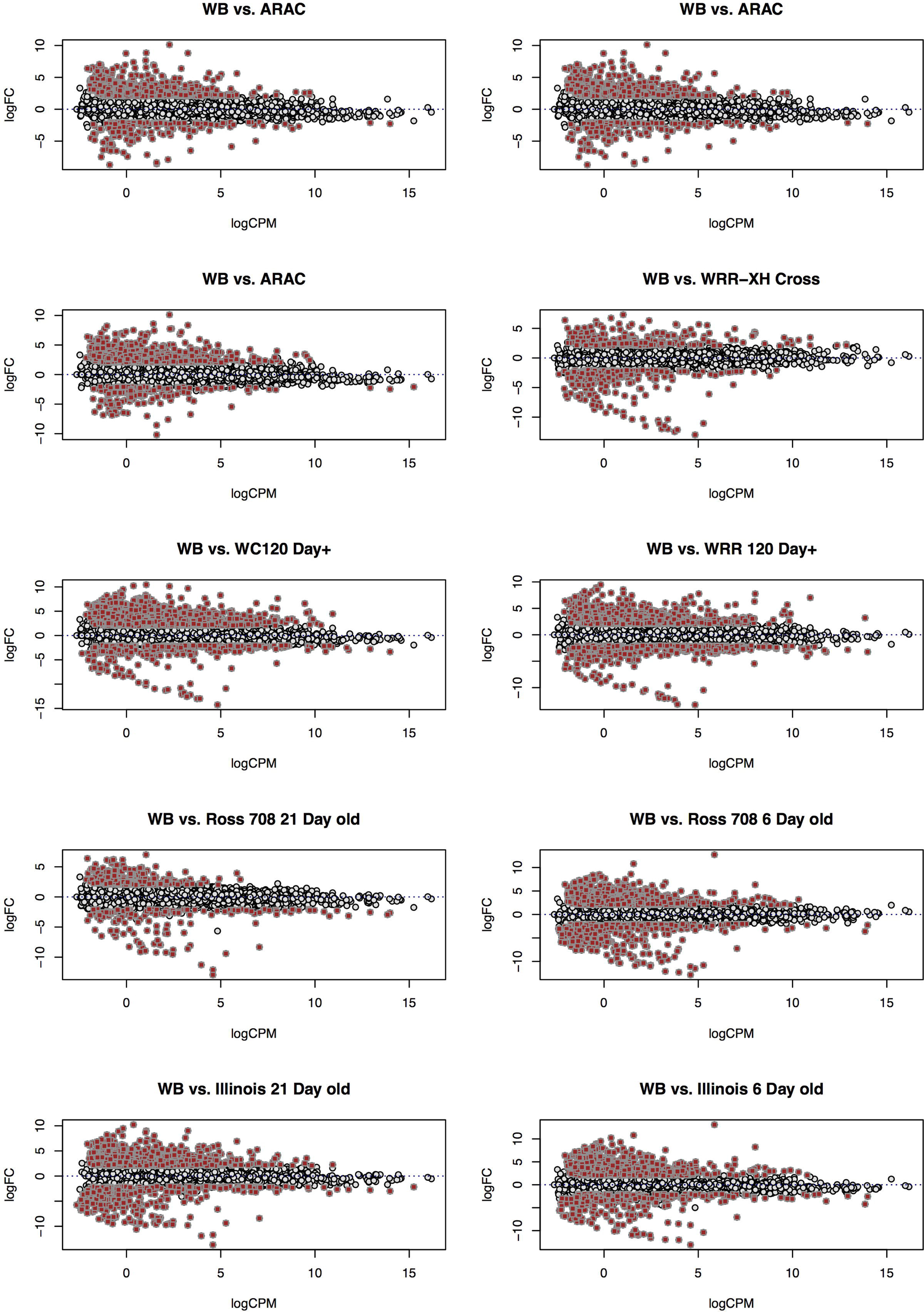
Mean-Average plots for analysis of differentially expressed genes for each pairwise comparison performed against the woody breast sample set. Points in red show the genes that were expressed at <-2 logFC or > 2 logFC, with a FDR <0.05.

**Table 2:**
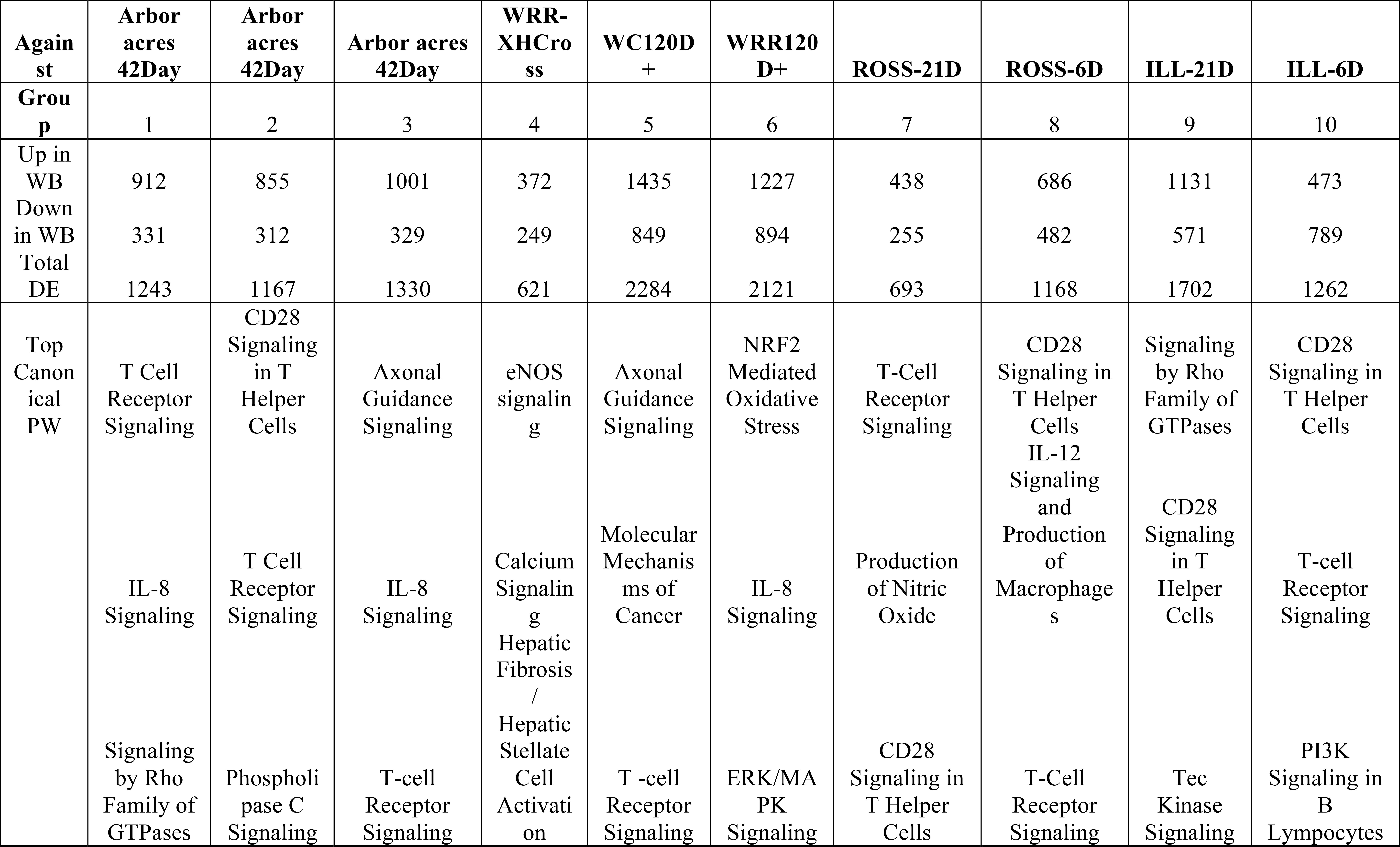
Summary of results from pairwise contrasts performed among RNAseq data using edgeR. The table shows the information on differentially expressed genes, summary of pathway analysis using IPA, upstream regulators for each dataset, and top disease terms.

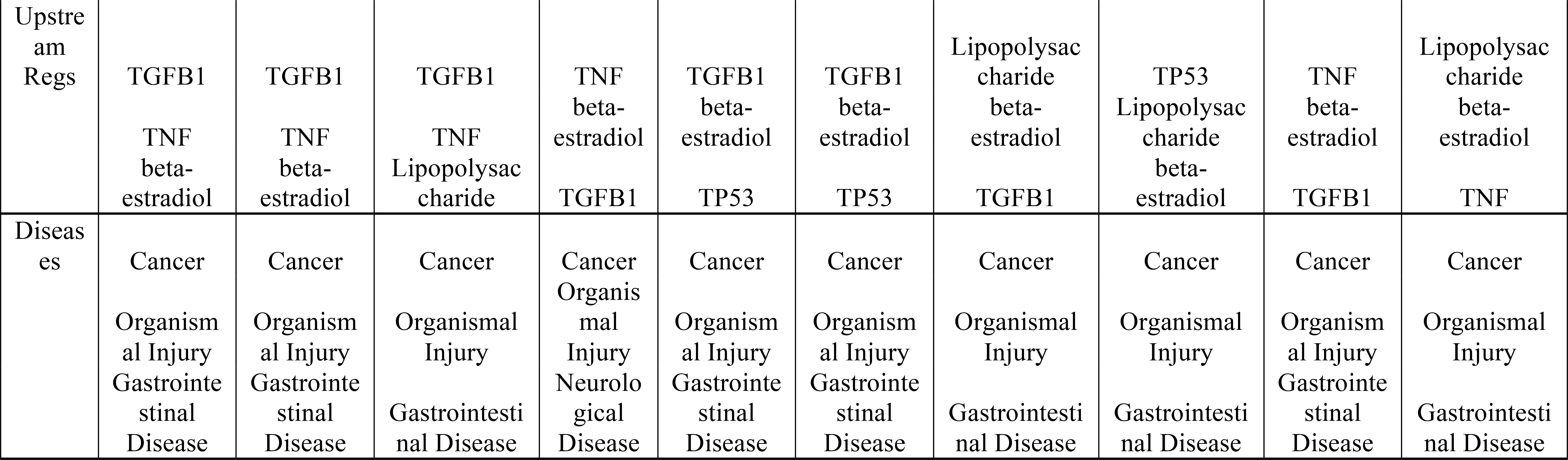

Comparing WB to Ross 708 (6 and 21-days old) showed more significant differences at six days than at 21-days, with main pathway terms being CD28 Signaling in T-helper Cells and T-cell receptor signaling. Lipopolysaccharide and beta-estradiol were shared, upstream regulators. Unlike the Ross strain, the Illinois 6 and 21-day old birds were distinct in their global expression profile, with more significant differences to WB compared to the younger Ross strain birds. This observation would fit the known biology of the Illinois strain, which is a broiler line with a performance profile from the 1950’s. However, pathway terms for these datasets still included CD28 signaling in T-helper Cells and T-cell receptor signaling, which were shared with the Ross strain. Both WC120+ and WRR120+ slow-growth breeds were most different from WB samples in the extent of differential expression, and with no overlap in the top three pathway terms.

Pathway analysis of genes differentially expressed in slow growth varieties also yielded pathway terms that were not shared with other comparisons. Pathway analyses yielded hepatic fibrosis/hepatic satellite cell activation, calcium signaling, eNOS signaling, molecular mechanisms of cancer, NRF2 mediated oxidative stress, ERK/MAPK signaling, signaling by Rho family GTPases, Tec kinase signaling, and PI3K signaling in B lymphocytes as the top terms in ARAC (3 libraries), WRR-XH cross, WC120D+, WRR120D+, Ross 6D and 21D, and Illinois 6D and 21D respectively‥ Interestingly, despite the differences in the top canonical pathways identified when comparing WB gene expression to that of slow growth varieties rather than fast-growth varieties, the upstream regulators suggested by this differential gene expression include the same terms. Diseases identified by the differential gene expression of each comparison evaluated included the same terms: Cancer, organismal injury, and gastrointestinal disease, with only one comparison indicating neurological disease. Finally, The frequent occurrence of T-cell receptor signaling and IL-8 Signaling suggest that molecules activated in these pathways may be suitable as biomarkers for detecting WB.

### Multisample Comparison Analysis

The multisample comparison analysis allows identification of similarities and trends occurring across multiple datasets, specifically, identification of functions overrepresented across datasets. Pathways are considered significant if a higher number of molecules associated with a pathway are expressed than expected by chance. Based on the pathways with highest −log(P-values) and activation Z-scores, the top shared canonical pathways were T-cell receptor signaling, CD28 signaling in T-helper cells, and signaling by Rho family of GTPases. The top 50 pathway and disease terms are shown in Table 3. The top upstream regulators were TGFb1, TNF and TP53 and beta-estradiol. The comparison feature also generated a list of the top diseases and disorders; the top three disease terms that emerged from the consensus of the multisample comparison were Cancer, Abdominal Neoplasm, and Solid Malignant Tumor. None of these top 100 disease terms pointed to muscle myopathies or other musculoskeletal disorders. Finally, the top disease signaling pathways were Cancer, MAPK and the P53 pathways.

**Table 3:**
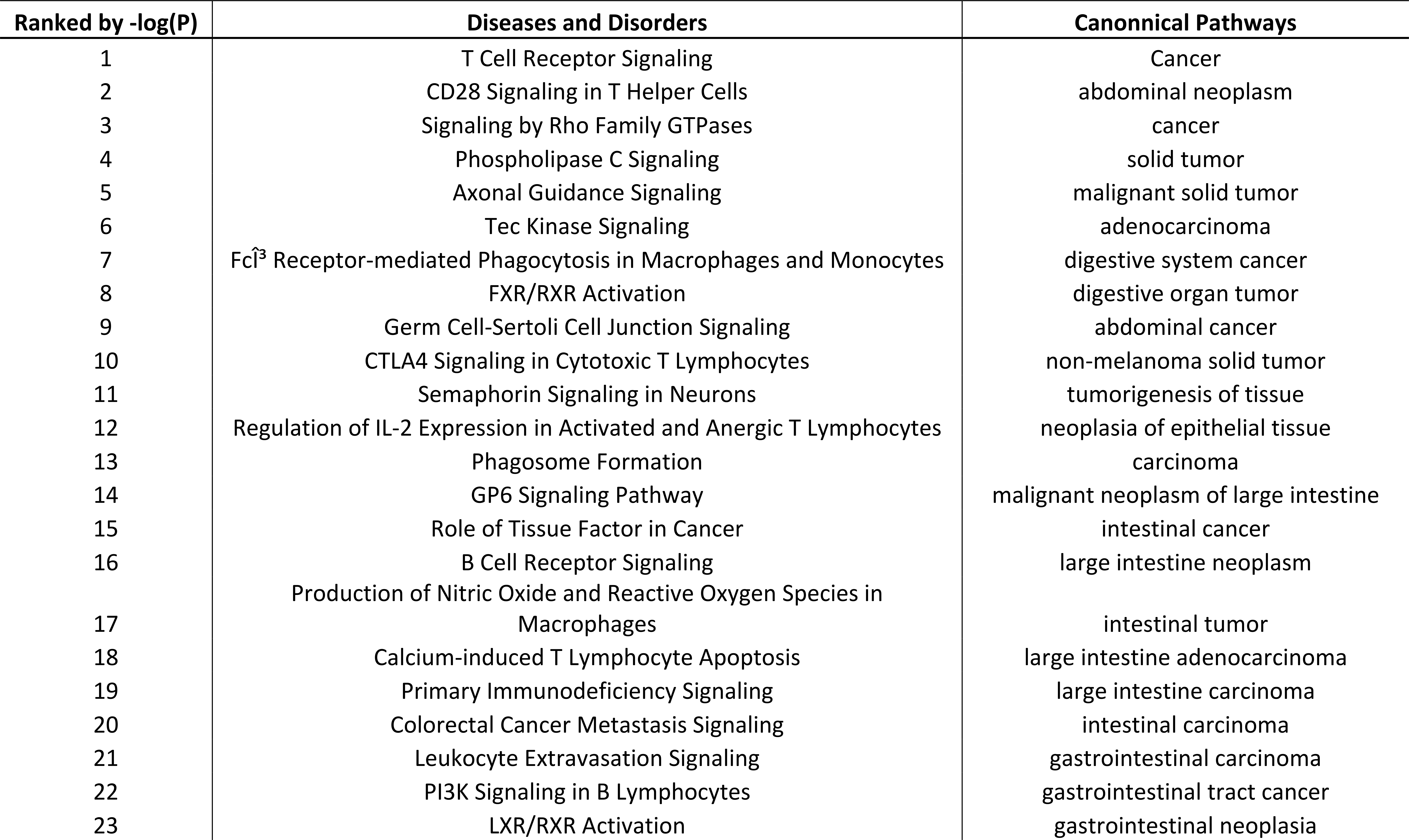
List of top 50 canonical pathway terms, and diseases and disorders identified from comparative analysis. In this analysis, multiple DEG datasets are compared to identify common pathway terms that are found more often than expected by chance.

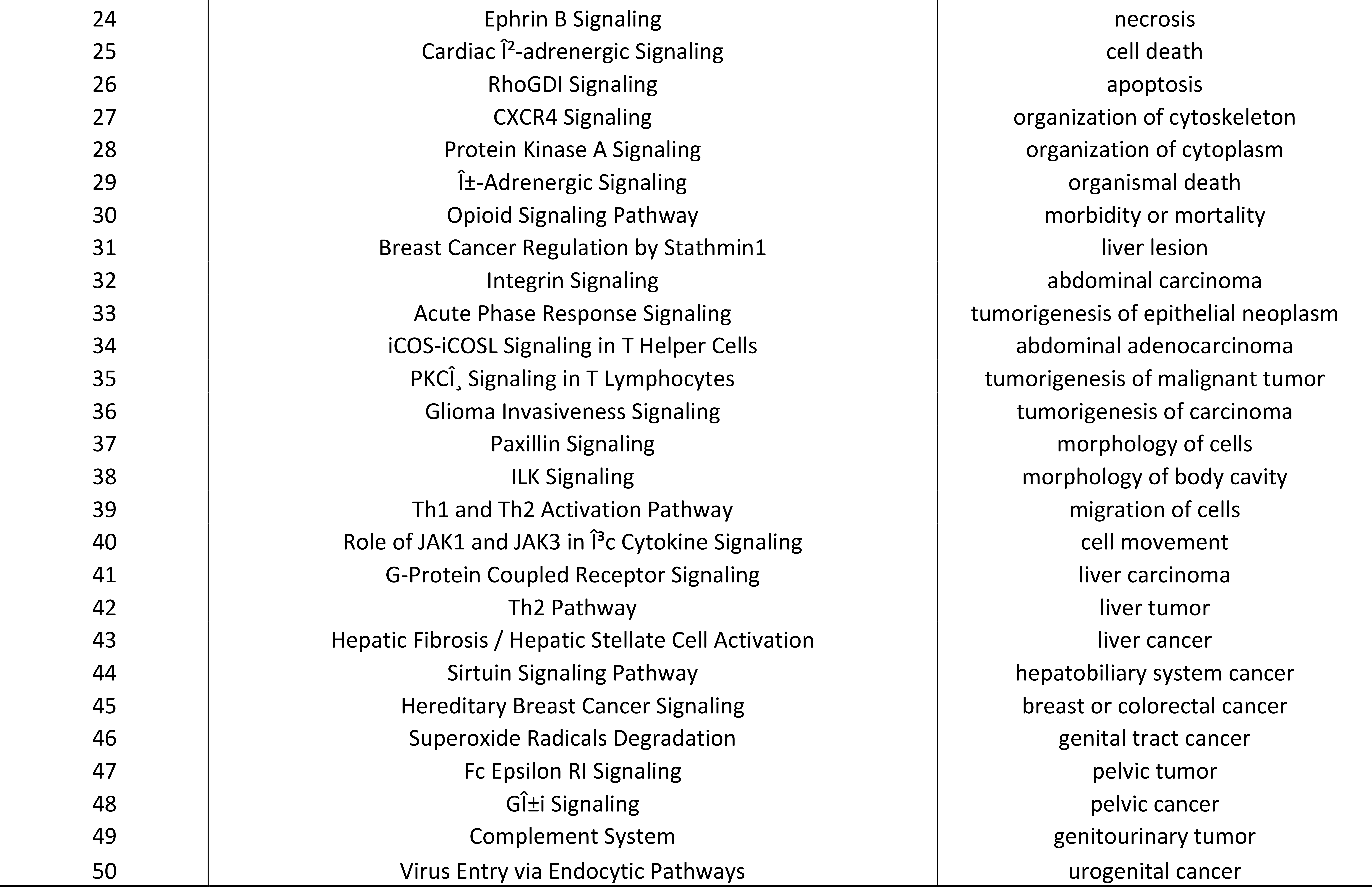

### Fast-Growth versus slow-growth differential expression

For this analysis, commercial broiler strains (ARAC, ROSS, WB) were included in the fast-growth commercial broiler (FGCB) group, and the WC120+ and WRR120+ strains were grouped into the slow-growth heritage broilers (SGHB) group. This particular analysis was designed to separate out those genes that are upregulated or downregulated in FGCB irrespective of age, with the supposition that genes associated with the onset and progression of WB in fast-growth strains would be found across age categories (6-days to 42-days). A total of 11,766 genes were expressed above a threshold (CPM=>2), of which 6406 were differentially expressed (FDR<0.05, LogFC < −0.05 and >0.05). Of the total differentially expressed genes, 6168 genes were significantly downregulated in FGCB. These differentially expressed genes were then analyzed in IPA to identify canonical pathways and diseases/disorders. The top three canonical pathways based on −log(P) values were Mitotic Roles of Polo-like Kinase, ILK Signaling, and ERK/MAPK signaling. As observed with other pairwise comparisons, the main disease terms were Cancer, Solid Malignant Tumors, and Gastrointestinal Disease.

### Variant discovery from RNAseq data

Variant calling and filtering of the datasets yielded average SNP calls of 79,539, and 83,902 for the FGCB and SGHB groups respectively. The individual variant calls were merged using GATK CombineVariants to generated a consolidated (merged) set of 709,959 and 279,339 high-quality SNPs in FGCB and SGHB respectively. This difference in total variants was driven mainly by differences in the number of samples included in each group – namely 25 and eight for FGCB and SGHB respectively. The multisample VCFs annotated with snpEff to identify effects of the variants and to categorize impacts yielded 395 and 158 high-impact variants in FGCB and SGHB respectively. Overall, the proportions of effected features and functional impacts were evenly matched (Table 4) except where noted. Modifier effects (changes outside coding regions) was the most frequent effect, which was higher in FGCB compared to SGHB. High-impact variants which signify impacts within the coding sequences (e.g., frameshift, stop gained, stop lost), were of equal proportion in both groups, whereas both moderate, and low-impact variants were more frequent in SGHB.

**Table 4:**
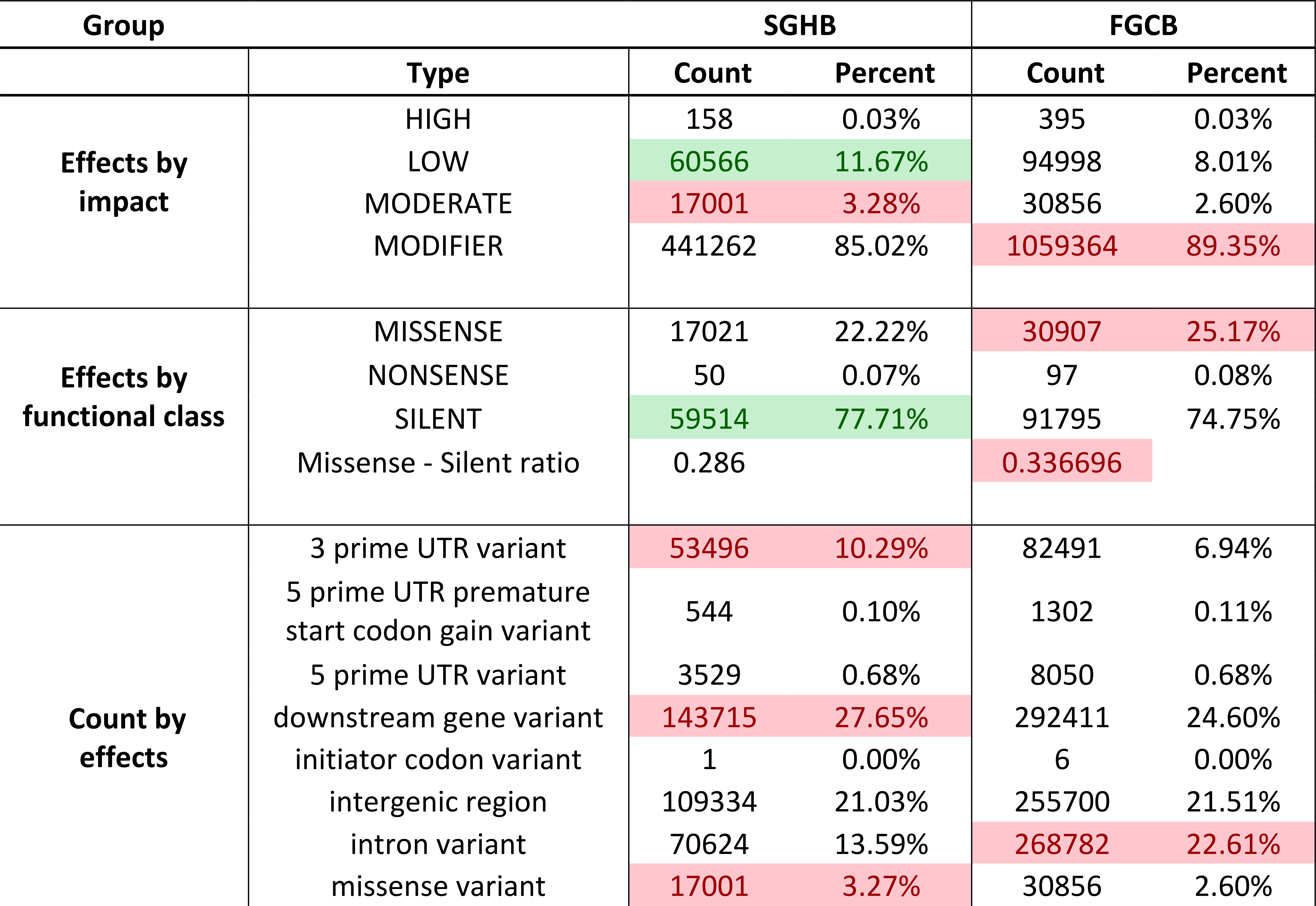
Comparison of variant effect prediction between the slow-growth heritage broilers (SGHB), and the fast-growth commercial broilers (FGCB). Results from snpEff based on high-quality SNP variants are shown. Colored boxes highlight notable differences in predicted effects for any category. Green colored boxes show elevated frequency of effects that are less likely to cause adverse impacts, whereas red shaded boxes show elevated frequency of phenotype-changing effects.

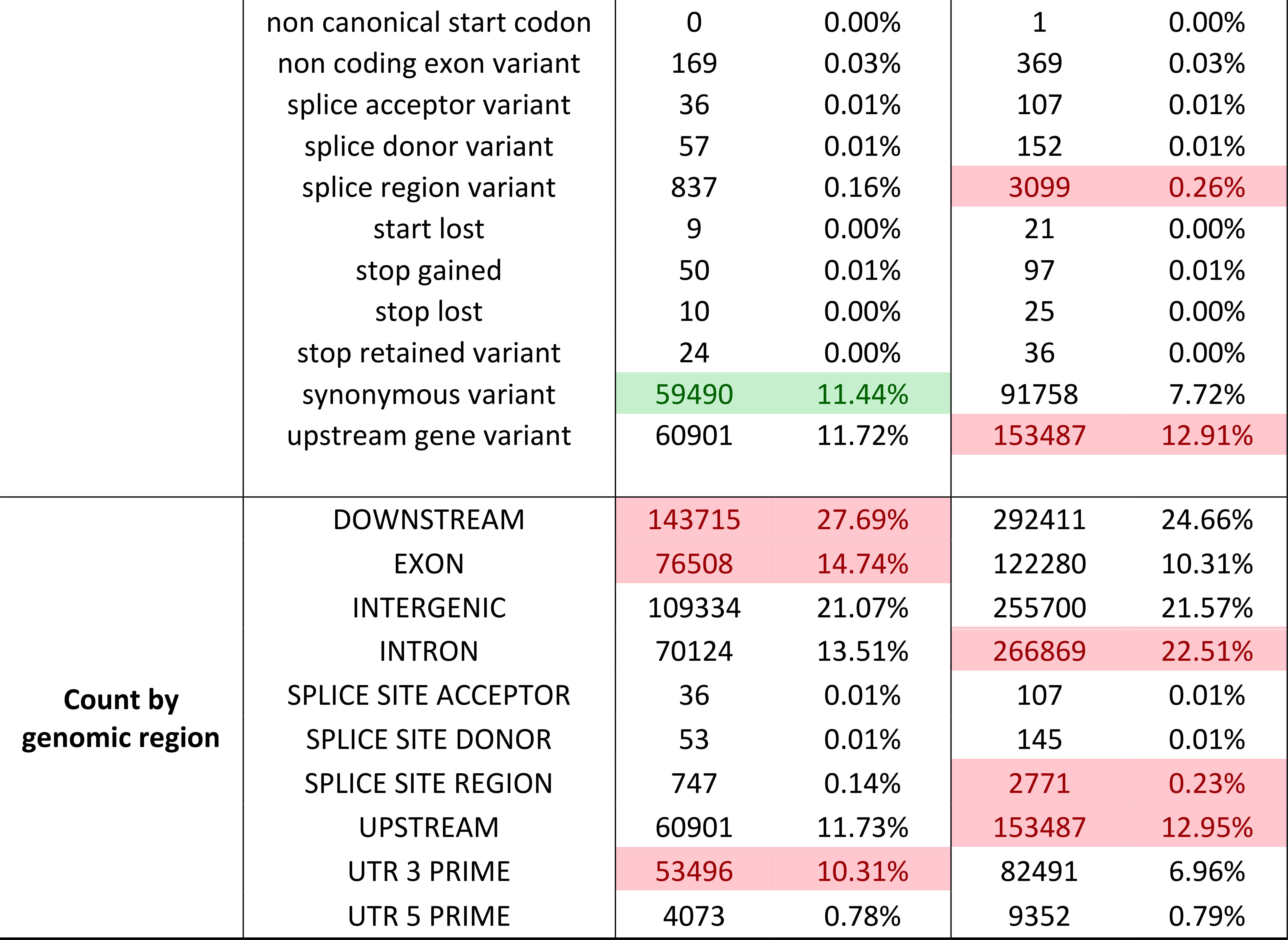

Of the total high-impact variants, 37 were shared among all three fast-growth breeds (ARAC, Ross, WB), whereas 35 were shared among the two slow-growth breeds (WC120+ and WRR120+). Of these 72 total high-impact variants, 14 were found in both the slow and fast-growth breeds, leaving 23 unique high-impact variants in FGCB, and 21 unique high-impact variants in SGHB (Table 5). The membership of both lists is rich in genes involved in cell signaling, cell proliferation, and cellular response to stress (including hypoxia). Particularly notable genes with high-impact variants unique to the FGCB group are SPEG, NPEPPS, and THYN1. These genes are involved in myocyte cytoskeletal development, linked to the cellular response to hypoxia, and associated with the induction of apoptosis, respectively. Notable genes with high-impact variants in the SGHB group were two myosin heavy chain genes (MYH1A, and MYH1B) and dystonin (DST). These genes are involved in motor activity and actin filament binding, and the assembly of collagen fibrils, respectively.

**Table 5:**
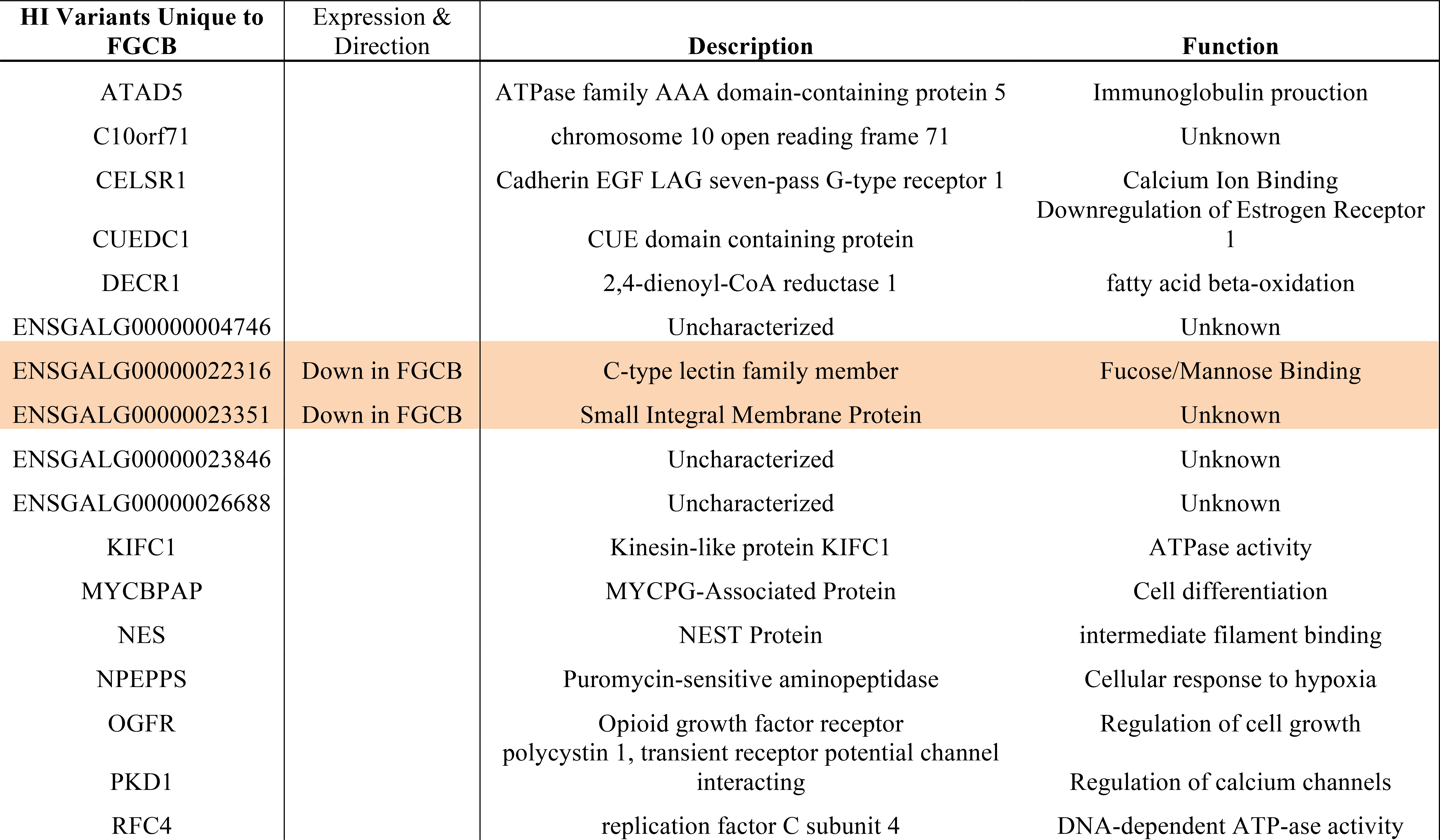
Lists of top high-impact variants unique to the fast-growth commercial broilers and slow-growth heritage broiler groups. Highlighted genes are those that were also significantly differentially expressed in comparison of Fast versus Slow groups. The directionality of regulation is also given for these differentially expressed genes.

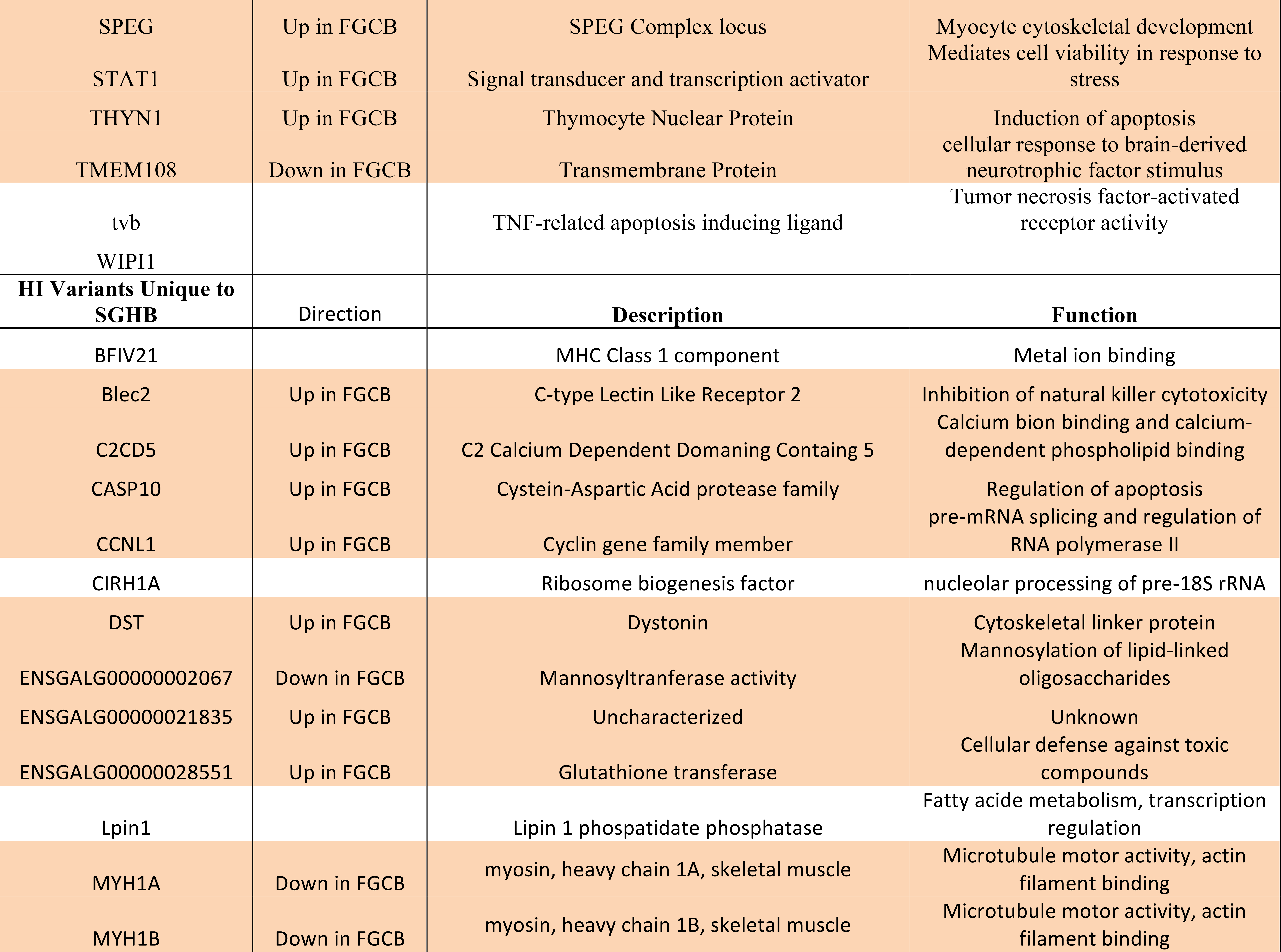

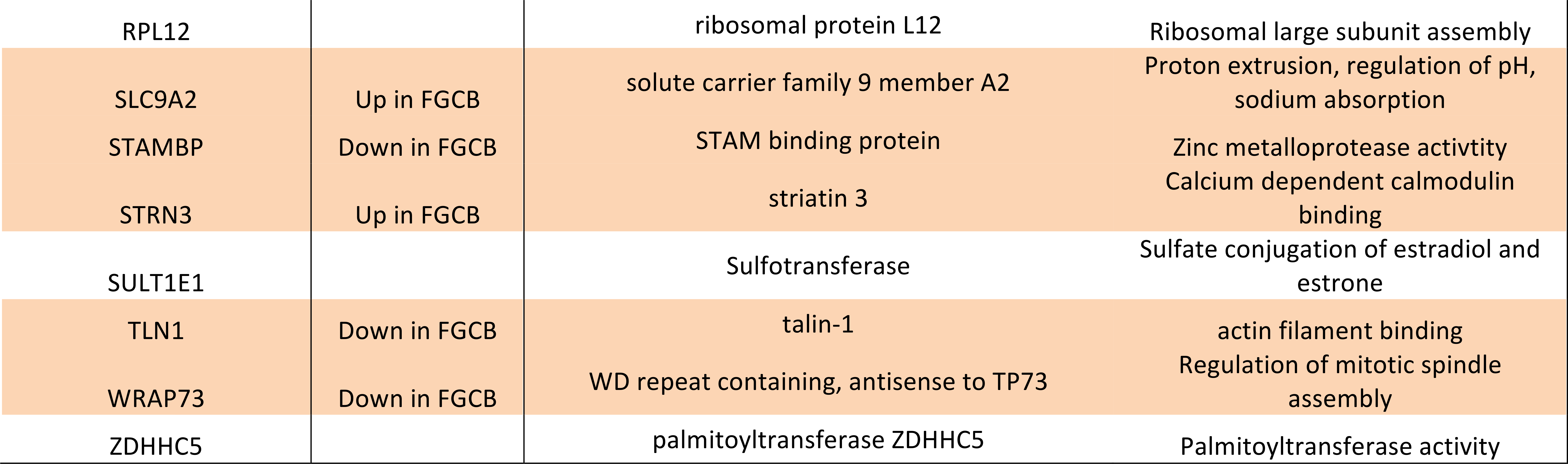

### Overlap of high expression and high-impact variants

Genes with high-impact variants in either FGCB or SGHB were cross-referenced against the list of significantly differentially expressed genes between the two groups. Twenty-three of the total 45 genes were also found to be significantly differentially expressed (Table 5). Genes that were up- or down-regulated in FGCB were found in both high-impact variant lists. Interestingly, 17 of the 21 genes with high-impact variants in SGHB were also significantly differentially expressed, suggesting both a mechanistic and functional role for these genes.

## Discussion

The hierarchical clustering of the pairwise differential expression analysis and the 100 top pathway terms shared across the 10 comparisons showed that there is a definite age based clustering pattern; all 42-day old broilers (ARAC) cluster together, and based on the Z-score and P-values, are more similar to WB+ tissue in their gene expression profiles and the pathways they activate. On the other hand, younger birds (6 and 21-days old) of slow and fast-growth breeds, and older birds of slow-growth breeds cluster together and separately from the 42-day old broilers. Interestingly, these results show that young broilers (Ross and Illinois 6 and 21-days old) and the slow-growth varieties (WC, WRR strains) appear to have a similar gene expression pattern, in contrast to 42-day old broilers. In summary, the comparison of pathways from multiple pairwise comparisons show that age, first, and then growth rate (broiler strain) are the primary functional differentiators of WB tissue. The differentiation of the 120+ day old slow-growth broilers and 6 and 21-day old fast-growth broilers is especially notable, as they show that a) molecular signatures associated with WB are unique to older, fast-growth broilers, and b) that 21-day old modern broilers (Ross 708) are less similar to 21-days old Illinois breed than to 42-day old commercial broilers. These two points suggest an age-dependent transcriptome dysregulation in WB, which progresses with age, somewhere between the first and third week of life. This conclusion is similar to that reached by Griffin *et al.* [19].

While gene expression and ontology analyses show which specific genes and molecular functions are involved in WB, the design of appropriate remediation strategies requires a better resolution of the similarity of WB to known diseases. A clear understanding of diseases and conditions explained by gene expression patterns is necessary to narrow down specific endogenous as well as environmental factors driving WB. Specifically, we wanted to answer whether WB tissue expression patterns point to muscle myopathies, or whether such patterns are indicative of other conditions. While genes essential in muscle growth and cell differentiation are up-regulated in WB tissue, the totality of expressed genes and pathways show little support for myopathy as the underlying condition.

One important cause for concern is the repeated occurrence of regulators and pathways that suggest neoplastic disorders. Upregulation of glycolysis, which was observed as the primary pathway classifier in the same-background comparison is considered a “near-universal property” of primary and metastatic cancers [39–42]. Oxidative stress and impaired glycolysis can both arise due to mitochondrial dysfunction. Multiple studies have confirmed the transcriptomic [9, 18] and metabolomic signatures of oxidative stress in WB [10], and have also suggested mitochondrial dysfunction [43]. Furthermore, genes essential to glycolysis, angiogenesis, and apoptosis (up-regulated in WB) are transcriptionally regulated in hypoxic conditions [44, 45], that are typical of tissue under oxidative stress.

The individual and comparison pathway analyses provided many of the same terms as being important among comparisons. The top pathways based on activation Z-score were T-cell receptor signaling, CD28 signaling in T-helper cells, and signaling by Rho family of GTPases. CD28 is involved in stimulation of T-cell activation and 23 molecules involved in this pathway were identified in the transcriptome data. CD28 signaling is involved in glucose metabolism, activation of T-cells, and costimulation of immune responses [46]. T-cell receptors bind to antigenic peptides presented by antigen presenting cells and are known to respond to various signal transduction pathways. They may be invoked to regulate cell proliferation, apoptosis, and cytotoxic killing [47, 48]. These processes may be activated in wooden breast in response to apoptosis and necrosis occurring in breast tissue. Finally, the Rho family of GTPases are involved in regulating various processes, including reorganization of the actin cytoskeleton in response to growth factors and cytokines [49–51]. These pathway terms are indicating abnormal expression patterns that together affect cell signaling, cytoskeletal organization, and inflammation have all been identified as features of WB. As it has been previously shown through histological and enzymatic assay that WS/WB does not have an infectious origin [4], these immune responses are likely directed against endogenous cell proliferation and apoptotic processes (resembling neoplasms).

The disease terms from the multigroup comparison analysis in IPA also found the same top conditions like those found in pairwise comparisons. Many of these terms also invoke the digestive system (intestine, colon, liver, abdomen, etc.), which is surprising considering all the analyses were based on differentially expressed genes in the pectoralis major tissue. While pathway analysis relies on over-representation or functional class scoring, they still rely on accurate annotations, cell-specific information, and well-described pathways for the accuracy of results [52, 53]. Therefore, while it is possible that other organs may be involved, validation of that question will depend on additional sampling and wet-lab based approaches. Other locations of organismal injury notwithstanding, these terms still suggest some important syndromes that can be considered very concerning. The major diseases and disorders explained by expression patterns also suggest abnormal cell proliferation and cell signaling mechanisms. The comparative analysis showed that over 80% of the top diseases identified by the pathway analyses indicate tumors, cancers, and neoplastic conditions. Even as some caution is necessary for interpreting pathway analysis for chicken datasets, due to the majority of pathways described being from mammalian models, it has repeatedly been shown that pathway signatures do predict disease outcomes accurately based on shared molecular features [54–57]. The reasons and basis for this similarity of WB to neoplastic disorders deserve further investigation.

Top canonical pathways emerging from the comparison between FGCB and SGHB groups indicated altered activity of multiple serine/threonine kinases including polo-like kinase (Plk), integrin-linked kinase (Ilk), and extracellular signal-regulated kinase/mitogen-activated protein kinase (Erk/Mapk). All three pathways are implicated in the regulation of the cell cycle and cell survival. Specifically, Plk is induced by mitogens and is most abundant during metaphase of mitosis with activities including chromosome segregation, centrosome maturation and spindle assembly [58, 59]. Plk also functions in other stages of mitosis including inactivation of the anaphase-promoting complex and regulation of nuclear envelope breakdown during prophase [58–60]. It has also been shown that depletion of Plk in cancer cells induces apoptosis and it is now considered a high potential target for intervention [58, 60]. Ilk impacts the cell cycle and survival through activation of critical signaling pathways and stimulation of downstream effector proteins, while Erk/Mapk is similarly involved by activating many growth factors, cytokines and transcription factors [61–64]. Erk/Mapk further promotes cell survival by phosphorylating and thus inhibiting the pro-apoptotic protein BAD (Bcl2 associated agonist of cell death) while also inducing the expression of cell survival genes [63]. Finally, of considerable interest is the ability of Ilk to anchor actin filaments to cell-matrix contact sites, regulating changes in cell shape, cell migration, cell adhesion, as well as the ability to phosphorylate myosin in smooth muscle cells [61, 62]. Each of these pathways plays an essential role in the regulation of cell proliferation, maintenance, and death which are all physiological activities that have been identified as perturbed in the WB condition and thus this analysis provides a focused framework for further investigating the underlying mechanisms of the condition.

### Molecular Basis of WB

In studies of WB, it has been observed that few live bird or carcass quality variables are accurately predictive of the presence of WB (Athrey et al., unpublished). For example, in replicate flocks of broilers of the same breed raised under identical conditions, no combination of rearing or dietary variables has been found to prevent WB occurrence [22, 65]. Furthermore, the severity of WB varies within the same flock under identical conditions [4]. Based on these observations, it appears likely that nutritional interventions, while perhaps useful for ameliorating severity, may be of limited utility in eliminating wooden breast. On the other hand, these data suggest a genetic basis underlying WB. This hypothesis has received recent support with the identification of QTL for WS [25]. While the relationships between WB and WS are not fully resolved, the co-occurrence of these two conditions makes it more likely that WB has a genetic basis.

In this study, we identified highly expressed genes that contained high-impact variants. This small subset of genes is involved in critical cell proliferation and signaling functions. The identification of genes with high-impact variants that are also differentially expressed point towards a genetic basis that links the changes at the DNA level to functional expression. Such DNA variants that are associated with expression differences are called Cis-acting regulatory variants and are known to explain a substantial amount of phenotypic variation, as well as having a role in disease etiology [66–68]. In this study, we identified a total of 44 high-impact variants, of which 21 were also significantly differentially expressed genes. While not all of these 21 genes may be directly affecting WB occurrence or severity, their expression patterns and the type of SNP modification make their involvement in WB highly probable. Particularly noteworthy genes identified in this analysis were MYH1A, MYH1B (high-impact variants in SGHB), the pair of which are myosin heavy chain genes. Both of these genes had ‘splice acceptor variants’ that may result in a different mature mRNA product and protein. These genes are members of a larger group of myosin genes which regulate development and function of avian skeletal muscle [69]. Interestingly, MYH1B was also identified as a candidate gene for white striping in a QTL mapping study by Pampouille *et al.* [25]. Another pair of notable genes with high-impact variants were DST (dystonin) and TMEM108 (Transmembrane protein). DST was modified in SGHB, whereas TMEM108 was modified in FGCB. DST is a cytoskeletal linker protein, which is involved in collagen trimerization and formation of scar tissue following injury [70]. DST is known to interact with TMEM108, a gene that regulates the stability of microtubules, by recruiting TMEM108 for the transport of endosomal vesicles [71]. Mutations in the DST gene have been identified as being responsible for hereditary neuropathy and dystonia (abnormal muscle tone) in mouse models [72]. The high-impact variants and differential expression of these genes may be associated with the rigidity, collagen content, and microscopic features observed in WB.

While the short list of genes identified in this study have a high likelihood of being explanatory of WB, and even as potential diagnostic biomarkers for the condition, we stop short of calling these candidate genes; population-level analyses such as association testing (GWAS) would be necessary to confirm if variants in these genes are causative of WB. However, these findings do lend support for a polygenic basis for WB, but perhaps occurring in conjunction with regulatory mechanisms that are yet to be identified and confirmed. Whether a putative genetic basis for wooden breast can be traced to linked loci under selection for growth traits or is a result of *de novo* mutations and structural variants, remains to be established. The high frequency of WB within flocks is suggestive of underlying causes that are not highly variable among individuals of a flock. Any variation in severity could, therefore, be a result of particular genotypes, and the resulting allele-specific growth traits and nutritional interactions. Such a pattern is consistent with multi-genic traits, alleles for which may segregate in populations as heterozygotes, and the occurrence and severity of the condition may be driven by allele-specific expression patterns [73, 74]. De novo mutations would also explain a *similar* pattern.

The results from our multisample comparison analysis show that WB has an age-dependent expression pattern, with molecular signatures and phenotypic markers becoming more evident in older birds. Such a phenomenon, called age-dependent penetrance, has frequently been observed in various heritable diseases. Recent studies have shown that some features of WB are observable as early as two weeks of age [19, 20], and therefore would explain our observation of 21-day old Ross birds being more similar to the older WB affected the group. The inherited human neurodegenerative disorder, Huntington’s Disease, is known to manifest in middle to later life, and gene expression studies show age-dependent expression and dysregulation of various signaling genes [75]. Age-dependent disorders may also include heritable genetic mechanisms such *de novo* mutations, or as somatic mutations [76–78] that may affect genome organization, or repair mechanisms and increase the penetrance of diseases in later life [79, 80]. Further investigations of genome organization, the frequency of *de novo* mutations, or breakdown of repair mechanisms in fast-growth broilers are necessary to illuminate whether and how these processes may be important in WB.

## Conclusions

Our study used transcriptomic datasets to compare pectoralis tissue from commercial broilers with wooden breast against multiple genotypic backgrounds, and confirmed the previously reported molecular signatures in addition to previously unreported molecules and pathways. The comparison of tissue from fast-growth genetic backgrounds to those from slow-growth genetic backgrounds and different age classes suggests a genetic basis for WB that elicits age-dependent expression patterns in fast-growth broiler strains. The functional analyses of pathways from comparative data suggest that WB is a potentially polygenic, complex syndrome, with molecular similarities to neoplastic disorders. Through analysis of high-impact variants among the studied breeds, we identified a short list of genes with high-impact variants that are also significantly differentially expressed, suggesting Cis-Regulatory processes involving essential developmental and cytoskeletal genes. This result underscores the need for more in-depth analyses to investigate the role of these genes, basis of these disease pathways and similarities to complex disorders.

## Ethical Approval statement

The animal studies and euthanasia were performed according to experimental protocols (Animal Use Protocol) approved by the Texas A&M University’s Institute for Animal Care and Use Committee (IACUC 2016-0065).

## Acknowledgments

We thank Jason Lee and Christine Alvarado for assistance with tissue sampling, and discussion of findings. This study was funded through start-up funding to the Athrey lab from Texas A&M AgriLife Research and Texas A&M University.

## Author Contributions

GA conceived, designed experiment design and carried out tissue sampling. SP performed wet lab work in preparation for sequencing. SP, TW, and GA conducted analyses, interpreted results, and generated figures. All authors reviewed the manuscript.

